# Kin28 depletion increases association of TFIID subunits Taf1 and Taf4 with promoters in *Saccharomyces cerevisiae*

**DOI:** 10.1101/634824

**Authors:** Elisabeth R. Knoll, Z. Iris Zhu, Debasish Sarkar, David Landsman, Randall H. Morse

## Abstract

In eukaryotes, transcription of mRNA-encoding genes by RNA polymerase II (Pol II) begins with assembly of the pre-initiation complex (PIC), comprising Pol II and the general transcription factors. Although the pathway of PIC assembly is well established, the mechanism of assembly and the dynamics of PIC components are not fully understood. For example, only recently has it been shown in yeast that the Mediator complex, which assists in pre-initiation complex formation at promoters of essentially all genes transcribed by Pol II, normally occupies promoters only transiently. This was inferred from studies showing that inhibiting Pol II promoter escape by depleting or inactivating Kin28 resulted in increased promoter occupancy by Mediator, as measured by chromatin immunoprecipitation (ChIP). Here we show that two subunits of TFIID, Taf1 and Taf4, similarly show increased occupancy as measured by ChIP upon depletion or inactivation of Kin28. In contrast, TBP occupancy is unaffected by depletion of Kin28, thus revealing an uncoupling of Taf and TBP occupancy during the transcription cycle. Increased Taf1 occupancy upon Kin28 depletion is suppressed by depletion of TBP, while depletion of TBP in the presence of Kin28 has little effect on Taf1 occupancy. Taf1 occupancy relative to TBP is higher at TFIID-dominated promoters and promoters having consensus TATA elements than at SAGA-dominated promoters and promoters lacking consensus TATA elements, consistent with prior work, and the increase in Taf occupancy upon depletion of Kin28 is more pronounced at TFIID-dominated promoters. Our results support the suggestion, based on recent structural studies, that TFIID may not remain bound to gene promoters through the transcription initiation cycle.

**Author Summary:** Transcription of mRNA-encoding genes by RNA polymerase II (Pol II) begins when the pre-initiation complex, a large complex comprising Pol II and several general transcription factors, including the TATA-binding protein (TBP)-containing TFIID complex, assembles at gene promoters. Although the major steps in the pathway of PIC assembly have been identified, the mechanism of assembly *in vivo* and the dynamics of PIC components are not fully understood. In this work we have used a yeast strain that is engineered to allow inhibition of promoter escape by Pol II by administration of a chemical, in order to “freeze” the assembled PIC and thus determine whether this condition increases the promoter occupancy of TBP and two TBP-associated factors (Tafs) that are components of TFIID. This approach was used recently to demonstrate that the Mediator complex, which facilitates PIC assembly, normally binds only transiently to gene promoters. We find that Tafs, like Mediator, show increased occupancy when Pol II promoter escape is inhibited, whereas TBP binding is constant. These results imply that binding of TBP and Tafs is uncoupled during the transcription cycle, and that Taf occupancy is at least partially interrupted upon Pol II promoter escape.

## Introduction

Transcription of mRNA genes in eukaryotes entails the formation of a pre-initiation complex (PIC) that includes the general transcription factors and Pol II. Although the paradigm for PIC formation, Pol II initiation and promoter escape is well established [1], major mechanistic questions remain. For example, although one of the earliest steps in PIC formation is promoter binding by TBP, how this occurs is uncertain. In yeast, TBP can be delivered to promoters by either the SAGA complex or TFIID [2, 3], and yeast promoters have been categorized as SAGA-dominated or TFIID-dominated based on relative occupancy by TFIID-specific subunits such as Taf1 and their response to mutations in SAGA or TFIID components [4–6]. However, transcription of genes in both categories depends on both SAGA and TFIID components, and whether distinct mechanisms of PIC formation and transcription initiation operate at the two classes is unknown [4, 7–9]. Further complicating the picture, recent work suggests that Tafs (i.e., TBP-associated factors that are subunits of TFIID) may function in a step occurring post-initiation in the transcription cycle [10], while structural studies indicate major changes in TFIID configuration during and after TFIID recruitment and PIC assembly [11].

Another component critical to assembly of the PIC is the Mediator complex. Mediator is a multiprotein complex that is conserved across eukaryotes and is important for transcription of essentially all genes transcribed by Pol II [12]. Insight into Mediator recruitment and dynamics has been gained by combination of genome-wide localization experiments, utilizing ChIP-chip (chromatin immunoprecipitation followed by microarray analysis), ChIP-seq (ChIP followed by high throughput sequencing), and ChEC-seq (chromosome endogenous cleavage followed by high throughput sequencing), together with genetic manipulations in yeast [13–18]. These studies showed that although Mediator could be detected at upstream activating sequences (UAS’s) of many transcriptionally active genes in yeast, there were also many active genes at which little or no Mediator association was observed by ChIP, consistent with early studies using ChIP followed by qPCR at a more limited set of genes [19, 20]. The puzzle presented by these results was resolved by the discovery that prevention of promoter escape by Pol II resulted in increased Mediator ChIP signal at promoters genome-wide [15, 18]. These findings indicated that Mediator association with promoters is normally transient, with dissociation occurring rapidly upon Pol II escape.

Stabilization of Mediator at gene promoters was accomplished by inactivation or depletion of Kin28, a kinase that is a subunit of TFIIH that phosphorylates the carboxy-terminal domain of the largest subunit of Pol II, thereby facilitating promoter escape [15, 18]. We recently showed that this stabilization depends on Pol II, as it is suppressed if Pol II is depleted using the anchor away technique [21]. The transient occupancy of promoter regions by Mediator during the transcription cycle raises the question as to whether other components of the transcription machinery are stably bound or, like Mediator, bind and rapidly dissociate upon Pol II promoter escape. Here we have used ChIP-seq to address this question with respect to TBP and Taf components of TFIID.

## Results

### Inactivation of an analog-sensitive mutant of Kin28 stabilizes promoter occupancy by Taf1

Mediator association with gene promoters is difficult to detect by ChIP in yeast under normal growth conditions, but yields a clear ChIP signal upon depletion or inactivation of Kin28 [15, 18]. Kin28 phosphorylates Ser5 of the YSPT**S**PS heptad repeat of the carboxy terminal domain (CTD) of Rpb1, the largest subunit of Pol II, facilitating its association with factors involved in transcriptional elongation and associated processes [22]. Inhibition of this phosphorylation by depletion or inactivation of Kin28 inhibits promoter escape by Pol II, and this evidently stabilizes Mediator association, thus implying that Mediator normally occupies active promoters only transiently [15, 18].

To test whether TFIID exhibits similar behavior, we monitored Taf1 association by ChIP in wild type yeast and yeast harboring an analog sensitive *kin28-as* mutation, in which Kin28 can be inactivated by administration of NaPP1 [23]. The *kin28-as* strain also harbored a *bur2*Δ mutation, to eliminate residual Ser5 phosphorylation by the Bur1/Bur2 complex [24]. ChIP against Taf1 followed by qPCR analysis revealed substantially increased Taf1 occupancy upon Kin28 inactivation at *RPL12A*, and modest increases in ChIP signal at the *PIK1* and *ARO3* promoters (Fig. 1A). No change in occupancy was apparent at the *TEF2* promoter, and a negative control, *SPS19*, showed no enrichment for Taf1 whether Kin28 was active or inactive. Taf1 occupancy did not differ between wild type yeast and *kin28-as bur2*Δ yeast, indicating that the *bur2* deletion did not affect Taf1 occupancy on its own.

**Figure 1.**
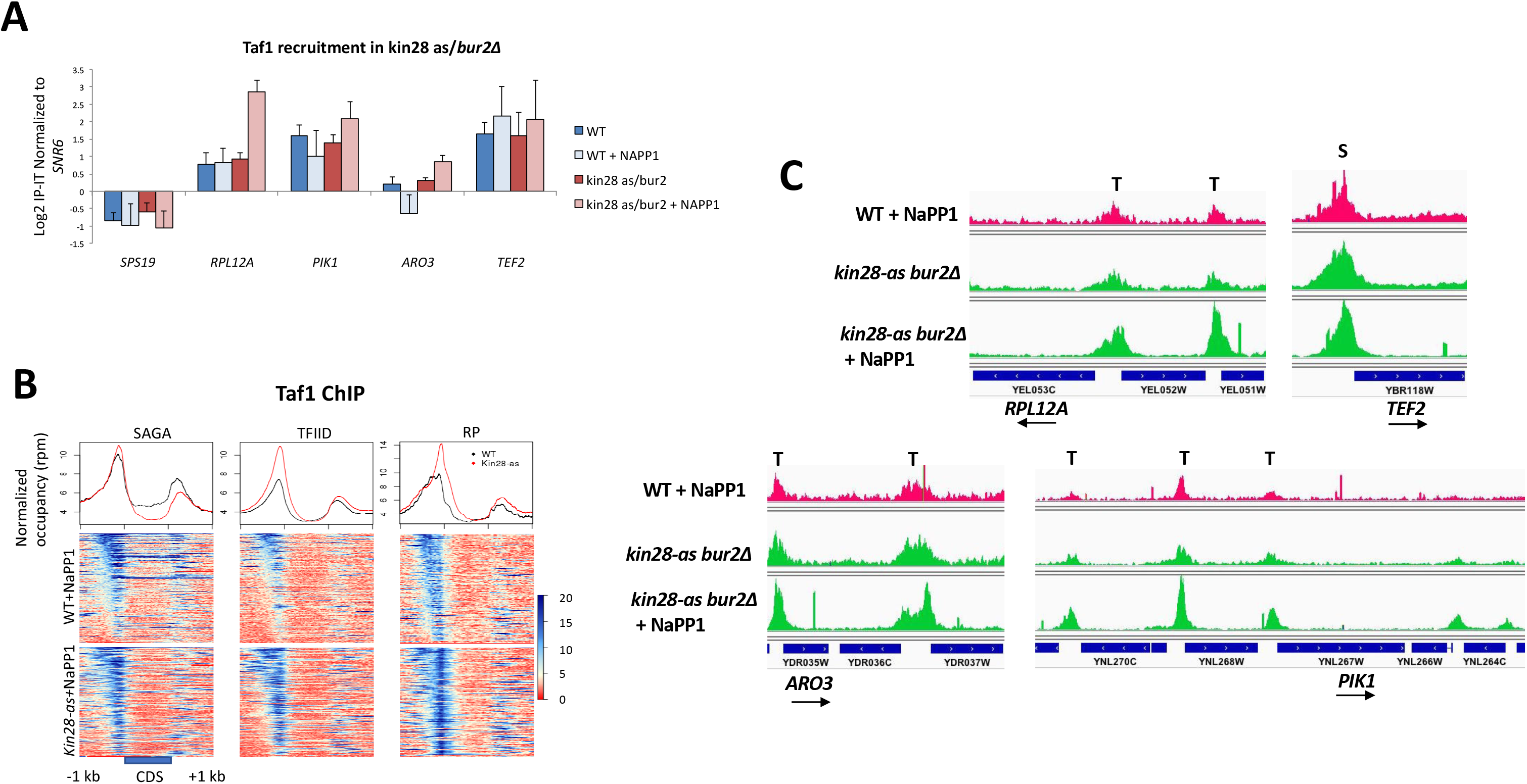
Increased occupancy of Taf1 at gene promoters upon inactivation of Kin28. (A) Normalized Taf1 occupancy at promoter regions of indicated genes was determined by ChIP followed by qPCR. ChIP was performed in wild type (WT) yeast (EPY4706) and *kin28-as bur2*Δ yeast (YFR912) grown in CSM-ura with or without 1 hr treatment with NaPP1 as indicated. Error bars reflect s.d.; n=3. (B) Normalized Taf1 occupancy in WT and *kin28-as bur2*Δ yeast after 1 hr treatment with NaPP1. Reads were mapped to all SAGA- and all TFIID-dominated genes, and to ribosomal protein (RP) genes. Genes were normalized for length and aligned by start and end of coding sequence (CDS) and sorted according to average signal intensity. Each horizontal line in the heat maps represents a gene, and the line graphs depict averages over all genes in the heat maps. (C) Browser scans showing normalized Taf1 occupancy in *kin28-as bur2*Δ yeast with and without NaPP1 treatment, and in WT cells treated with NaPP1. Peaks at SAGA-dominated and TFIID-dominated gene promoters are indicated by “S” and “T”, respectively.

To examine the effect of Kin28 inactivation on Taf1 occupancy on a genome-wide scale, we conducted ChIP followed by high throughput sequencing (ChIP-seq) against Taf1 in wild type yeast and in *kin28-as bur2*Δ yeast before and after treatment with NaPP1. Unlike Mediator subunits, Taf1 exhibits a clear ChIP signal near the transcription start site (TSS) of many genes under normal growth conditions. This signal increased upon inactivation of Kin28, particularly at TFIID-dominated genes, including both ribosomal protein (RP) genes and non-RP genes (Fig. 1B). ChIP-seq results for *RPL12A, TEF2, ARO3*, and *PIK1* were consistent with qPCR results, with all but *TEF2* showing increased occupancy of Taf1 upon Kin28 inactivation (Fig. 1C). Increased Taf1 occupancy was seen at other promoters as well, consistent with heat map results. These results suggest that Taf1 may, like Mediator, be stabilized at promoters when promoter escape by Pol II is inhibited, with the effect being strongest at TFIID-dominated promoters.

### Depletion of Kin28 stabilizes promoter occupancy by Taf1 and Taf4

To test further the effect of Kin28 on Taf occupancy, we used the anchor away method to deplete Kin28 from the nucleus [25]. This method employs a yeast strain in which the ribosomal protein Rpl13A has been modified by addition of a C-terminal FKBP12 tag, while the protein of interest (Kin28) is C-terminally tagged with the FRB fragment, which binds tightly to FKBP12 upon addition of rapamycin, resulting in the FRB-tagged protein being transported out of the nucleus during ribosomal protein processing. The anchor away yeast strain also harbors a *tor1-1* mutation, which abrogates the normal stress response induced by rapamycin [25]. Previous studies used this method to demonstrate increased association of Mediator with promoters, as monitored by ChIP, after depletion of Kin28 by 1 hr of rapamycin treatment [14, 17, 18].

We conducted ChIP-seq using antibodies against the TFIID-specific subunits Taf1 and Taf4 and observed a strong correlation (r^2^ = 0.95) between Taf1 and Taf4 occupancy over a wide range of peak intensities (Fig. 2A). Depletion of Kin28 by 1 hr of rapamycin treatment resulted in increased ChIP signal for both Taf1 and Taf4 at most promoters (Fig. 2B-D). Consistent with the results for the *kin28-as* mutant, the increase in both Taf1 and Taf4 occupancy was more pronounced at TFIID-dominated promoters than for SAGA-dominated promoters (Fig. 2D; p = 1.5 × 10^−12^ for Taf1 and p = 1.5 × 10^−5^ for Taf4 at SAGA vs. TFIID promoters (Mann Whitney U test)). This differential effect could be observed at individual promoters; compare the increase in signal upstream of the TFIID-dominated *NSG2/CUZ1* or *YAP1801* promoter to the SAGA-dominated *YGP1* or *MPC2* promoter (Fig. 2E).

**Figure 2.**
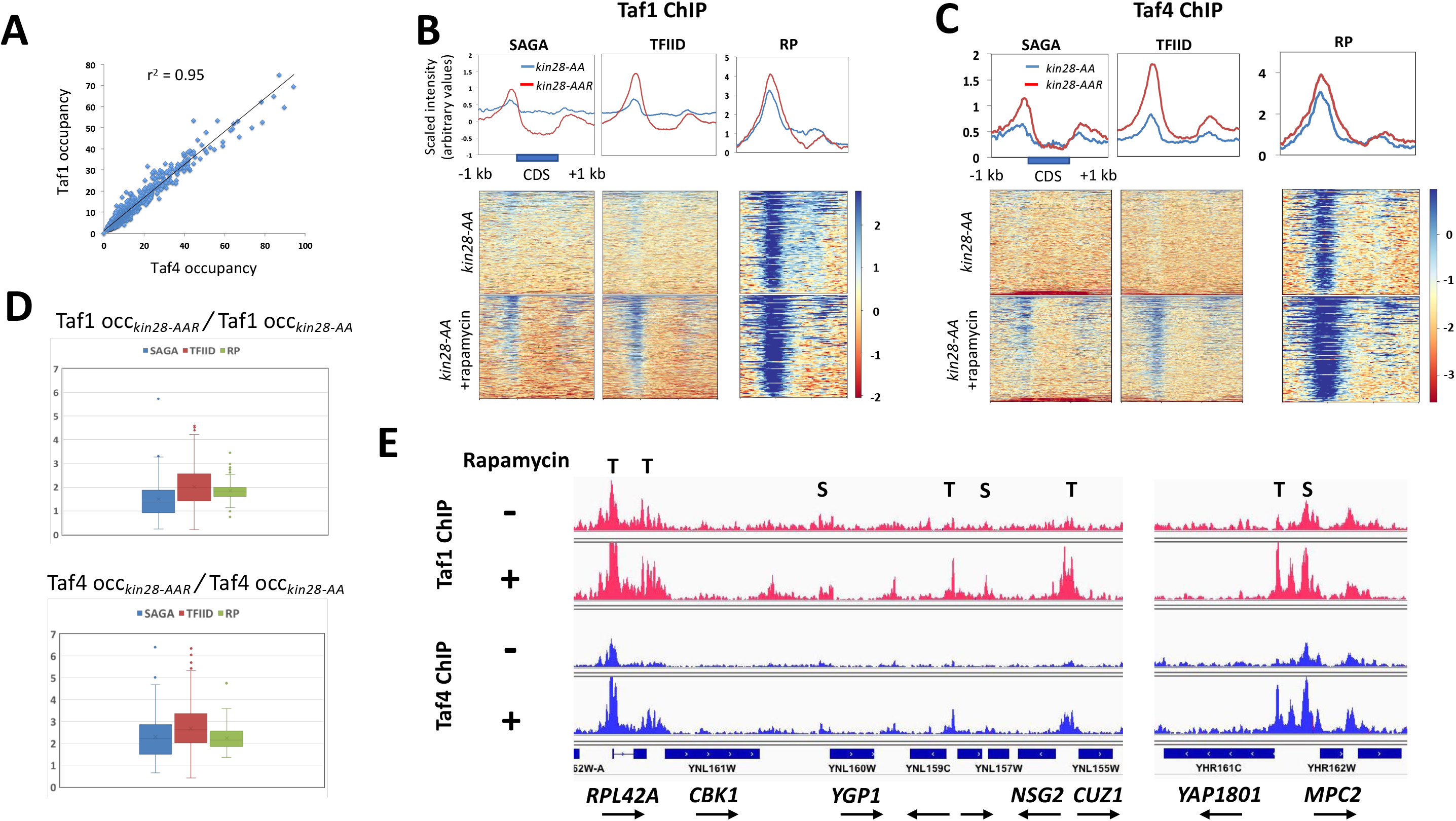
Increased occupancy of Taf1 and Taf4 at gene promoters upon depletion of Kin28. (A) Normalized occupancy of Taf1 plotted against Taf4 occupancy for the ~1000 genes (see Methods) most highly occupied by Pol II. (B-C) Normalized occupancy is depicted for Taf1 (B) and Taf4 (C) in *kin28-AA* yeast, without and with 1 hr rapamycin treatment, mapped to all SAGA-dominated, TFIID-dominated (excluding RP genes), and RP genes. Genes were normalized for length, aligned by coding sequence (CDS) start and stop, and sorted according to average signal intensity. The 19 SAGA-dominated genes at the bottom of the Taf4 heat maps were removed before calculating averages used in the line graphs, as these were almost all Ty1 elements that had higher intensity in the input control than in the Taf4 ChIP sample, and therefore yielded negative values in the heat map. Note that a different scale is shown for the RP gene line graphs than for SAGA- and TFIID-dominated genes. (D) Ratios of Taf1 (top) and Taf4 (bottom) occupancy in *kin28-AA* yeast in the presence and absence of rapamycin are shown in box and whisker plots for the ~1000 genes having highest occupancy by Pol II (see Methods), sorted into SAGA-dominated, non-RP TFIID-dominated, and RP genes. The boxes show the 2^nd^ and 3^rd^ quartiles, and the whiskers indicate the 1^st^ and 4^th^ quartiles; median values are indicated by the horizontal lines in the boxes separating 2^nd^ and 3^rd^ quartiles, and outliers are depicted as points above or below the whiskers. (E) Browser scans showing normalized Taf1 and Taf4 occupancy in *kin28-AA* yeast with and without rapamycin treatment. Peaks at SAGA-dominated and TFIID-dominated gene promoters are indicated by “S” and “T”, respectively.

We also examined TBP occupancy and found little effect of depleting Kin28 (Fig. 3A-B). TBP occupancy in the presence and absence of rapamycin in *kin28-AA* yeast showed essentially no change on average for TFIID-dominated genes, both RP and non-RP, and a slight decrease in occupancy at SAGA-dominated genes (Fig. 3B). Decreased ChIP signal for TBP following Kin28 depletion could be observed at select promoters in browser scans; Fig. 3C shows that TBP signal is decreased at *CDC19*, while being relatively unaffected at *CLN3* and the divergent *RBG1-FUN12* promoters. We noticed that anomalous behavior was also seen for Taf1 and Taf4 at some of the promoters showing decreased TBP signal following Kin28 depletion, with little or no increase in Taf1 signal when Kin28 was depleted; *CDC19* again illustrates this point (Fig. 3C). We do not have a mechanistic explanation for the decrease in TBP occupancy observed at select genes following Kin28 depletion. These results indicate that TBP occupancy, in contrast to Taf1 occupancy, is not increased by depletion of Kin28. Thus, while Taf1, like Mediator, binds transiently to most promoters and is stabilized under conditions that inhibit promoter escape by Pol II, TBP binding is stable and does not require continued occupancy by Mediator or the Taf proteins.

**Figure 3.**
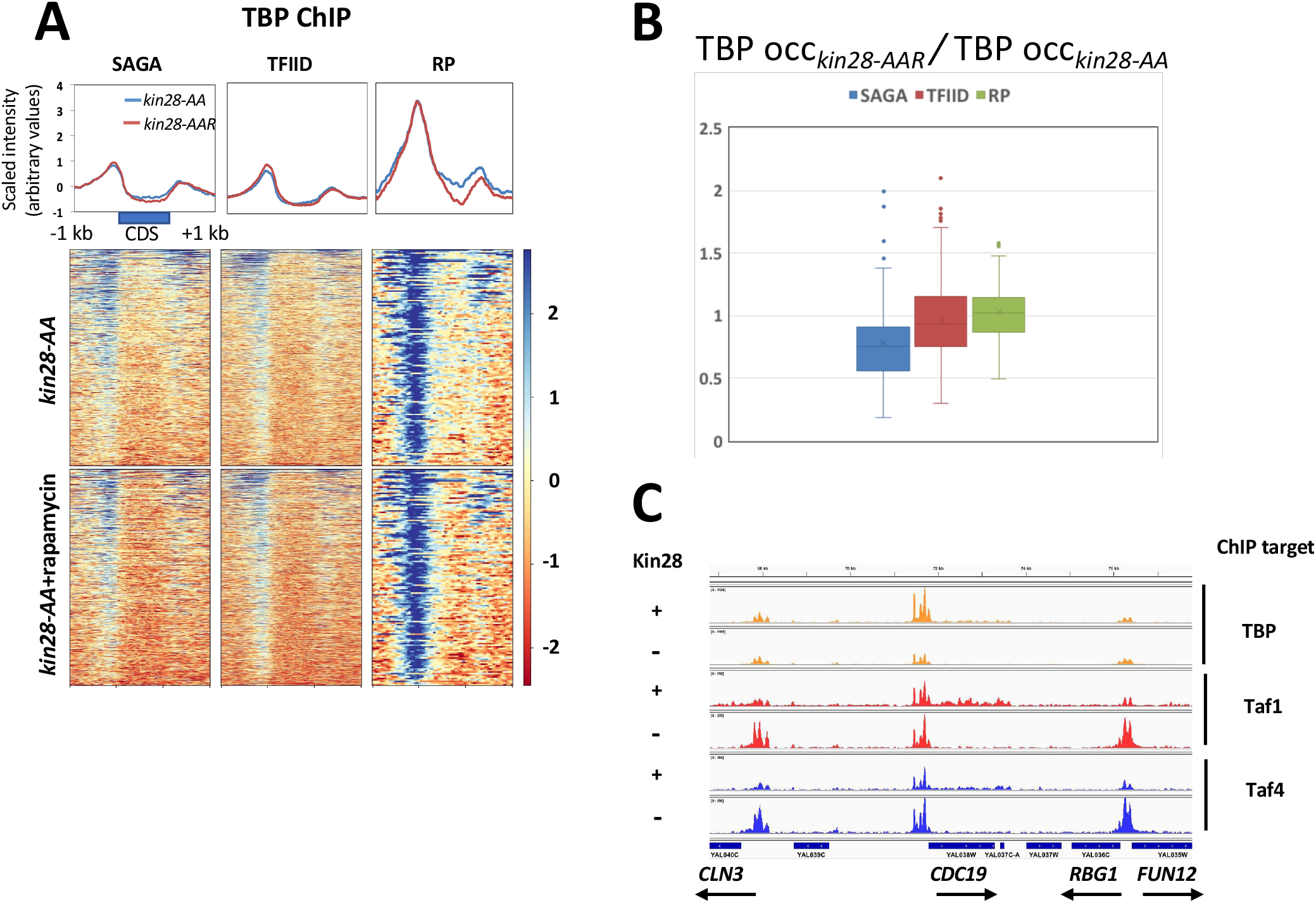
TBP occupancy is not affected by depletion of Kin28. (A) Normalized TBP occupancy in *kin28-AA* yeast, without and with 1 hr rapamycin treatment, mapped to all SAGA-dominated, non-RP TFIID-dominated, and RP genes. Genes were normalized for length, aligned by coding sequence (CDS) start and stop, and sorted according to average signal intensity. (B) Ratios of TBP occupancy in *kin28-AA* yeast in the presence and absence of rapamycin are shown in box and whisker plots, as in Figure 2, for the ~1000 genes having highest occupancy by Pol II (see Methods), sorted into SAGA-dominated, non-RP TFIID-dominated, and RP genes. (C) Browser scan showing normalized Taf1, Taf4, and TBP occupancy in *kin28-AA* yeast with and without rapamycin treatment.

### Increased occupancy by Taf1 upon Kin28 depletion is suppressed by TBP depletion

To test whether the increased association of Taf1 seen upon Kin28 depletion requires stable TBP occupancy, we performed ChIP-seq against Taf1 in *kin28-tbp-AA* yeast, in which rapamycin addition causes depletion of both Kin28 and TBP [21]. We found that Taf1 occupancy was essentially unchanged upon simultaneous depletion of Kin28 and TBP (Fig. 4A); marked effects on Mediator and Pol II association provide evidence for efficient depletion of both Kin28 and TBP [21], as does the loss of the artifactual, transcription-dependent signal at the 3’ ends of coding sequences (Fig. 4A) [16, 26]. Occupancy ratios for Taf1 and Taf4 in the presence and absence of rapamycin in *kin28-AA* yeast are increased by 1.5-3 fold at most genes upon depletion of Kin28, as discussed earlier (Fig. 4B). Depletion of TBP together with Kin28 abrogates this increased occupancy at most genes in all categories (Fig. 4B).

**Figure 4.**
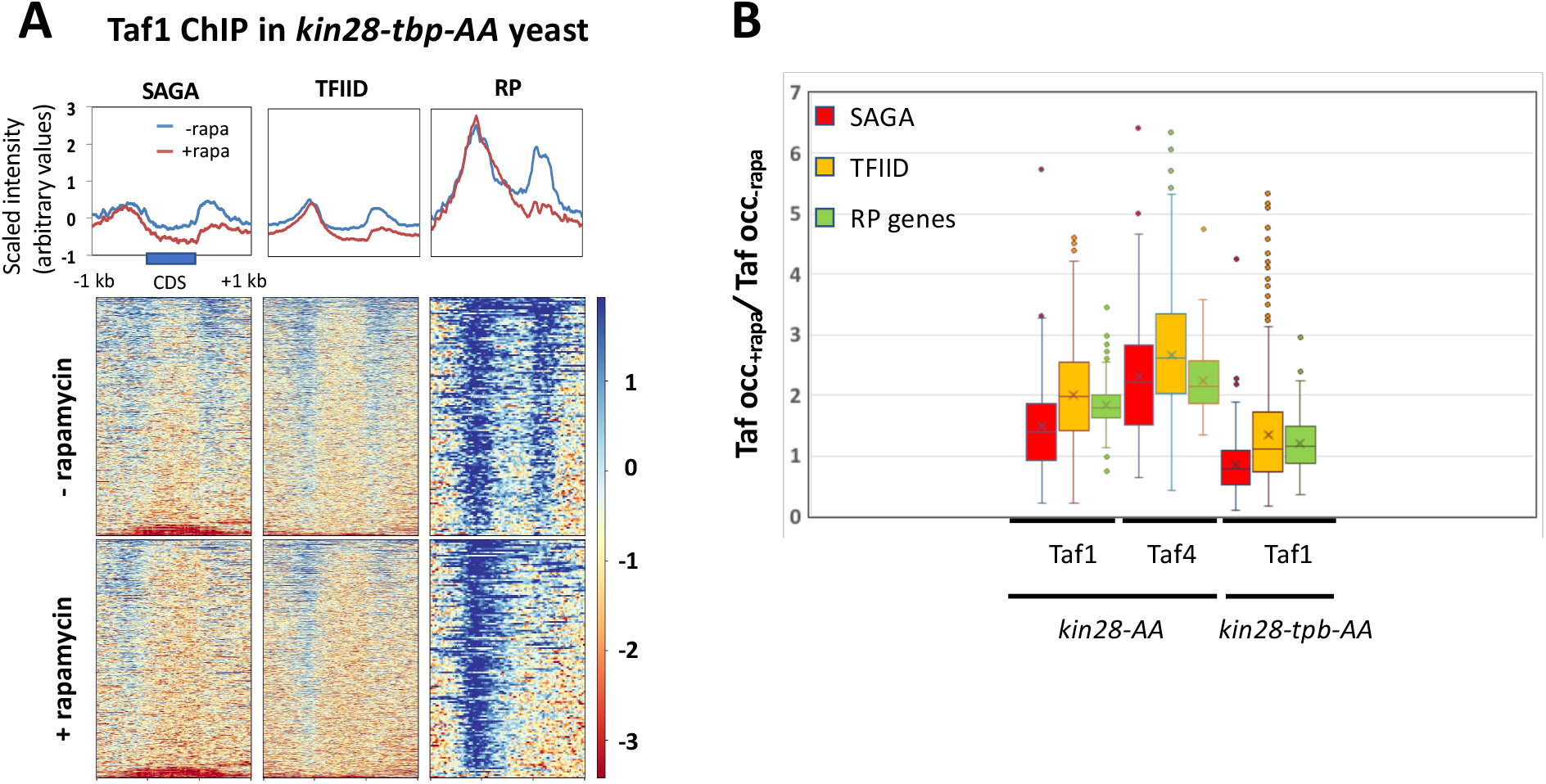
Depletion of TBP suppresses increased occupancy of Taf1 seen upon depletion of Kin28. (A) Normalized TBP occupancy in *kin28-tbp-AA* yeast, without and with 1 hr rapamycin treatment, mapped to all SAGA-dominated, non-RP TFIID-dominated, and RP genes. Genes were normalized for length, aligned by coding sequence (CDS) start and stop, and sorted according to average signal intensity. The 21 SAGA-dominated genes at the bottom of the heat maps were removed before calculating averages used in the line graphs, as these were almost Ty1 elements that had higher intensity in the input control than in the Taf1 ChIP sample, and therefore yielded negative values in the heat map. (B) Ratios of Taf1 and Taf4 occupancy in *kin28-AA* yeast in the presence and absence of rapamycin (same data as in Fig. 2) and of Taf1 occupancy in *kin28-tbp-AA* yeast in the presence and absence of rapamycin are shown in box and whisker plots, as in Figure 2, for the ~1000 genes having highest occupancy by Pol II (see Methods), sorted into SAGA-dominated, non-RP TFIID-dominated, and RP genes.

We noted that a substantial number of outliers among TFIID-dominated genes showed increased Taf1 association upon depletion of Kin28 and TBP. Analysis of this cohort (101 non-RP, TFIID-dominated genes showing > 2X increase in Taf1 association in *kin28-tbp-AA* yeast after rapamycin treatment) revealed that it was enriched for genes involved in mRNA binding (21 genes; corrected p-value 1.9 × 10^−11^) and related functions (e.g. organic cyclic compound binding), and for genes involved in ribosome biogenesis and related processes (53 genes; p-value 6.2 × 10^−34^) [27]. Furthermore, this gene set was enriched for promoters occupied by Abf1 but not by Rap1 (p-value 1.5 × 10^−4^ for Abf1 compared to p-value 0.09 for Rap1 for genes with association p-value < 0.005; hypergeometric test [28]). Possibly Taf1 is stabilized by factors associating with these promoters, including Abf1, such that TBP is not needed for its stabilization upon Kin28 depletion.

Based on these results, we conclude that stable association of TBP is required at most promoters for increased association of Taf1 upon depletion of Kin28.

### Taf1 occupancy does not require TBP

We recently reported investigations of the interdepencies among the PIC components TBP, Taf1, and Pol II for occupancy of promoter regions, using ChIP-seq before and after conditional depletion of these same components [21]. Examination of occupancy of PIC components at a subset of genes, dubbed “UAS genes”, that exhibit Mediator peaks at upstream activating sequences (UASs) in wild type yeast, revealed that depletion of TBP resulted in nearly complete loss of Pol II occupancy, but little change in occupancy by Taf1. These results are consistent with the generally accepted view that recruitment of Pol II strongly depends on TBP, and indicate that Taf components of TFIID can associate with promoters even in the absence of TBP. The latter observation is consistent with previous reports showing continued Taf1 association at several promoters after inactivation of a *tbp-ts* mutant [3, 6].

We were interested in determining whether promoters categorized as “TFIID-dominated” and “SAGA-dominated” differed in these interdependencies, as this could help address the question of whether such promoters differ mechanistically. We therefore re-examined our data to ascertain the effect of depletion of TBP on Taf1 occupancy at SAGA-dominated and TFIID-dominated genes. Examination of heat maps and line graphs indicated little change in Taf1 occupancy after depletion of TBP (Fig. 5A). TBP depletion was efficient as shown by ChIP-seq of TBP [21]. Quantitative comparison of Taf1 occupancy in the presence and absence of rapamycin in *tbp-AA* yeast revealed on average a decrease of about 30% at SAGA-dominated genes and no change at TFIID-dominated genes, including RP genes (Fig. 5B). Variable effects of TBP depletion on Taf1 occupancy were observed in browser scans, although most Taf1 peaks were unaffected by TBP depletion (Fig. 5C). We conclude that Taf1 occupancy does not generally depend on continued occupancy by TBP, and that SAGA-dominated genes show a mildly stronger dependence on TBP for normal levels of Taf1 association with promoters than do TFIID-dominated genes.

**Figure 5.**
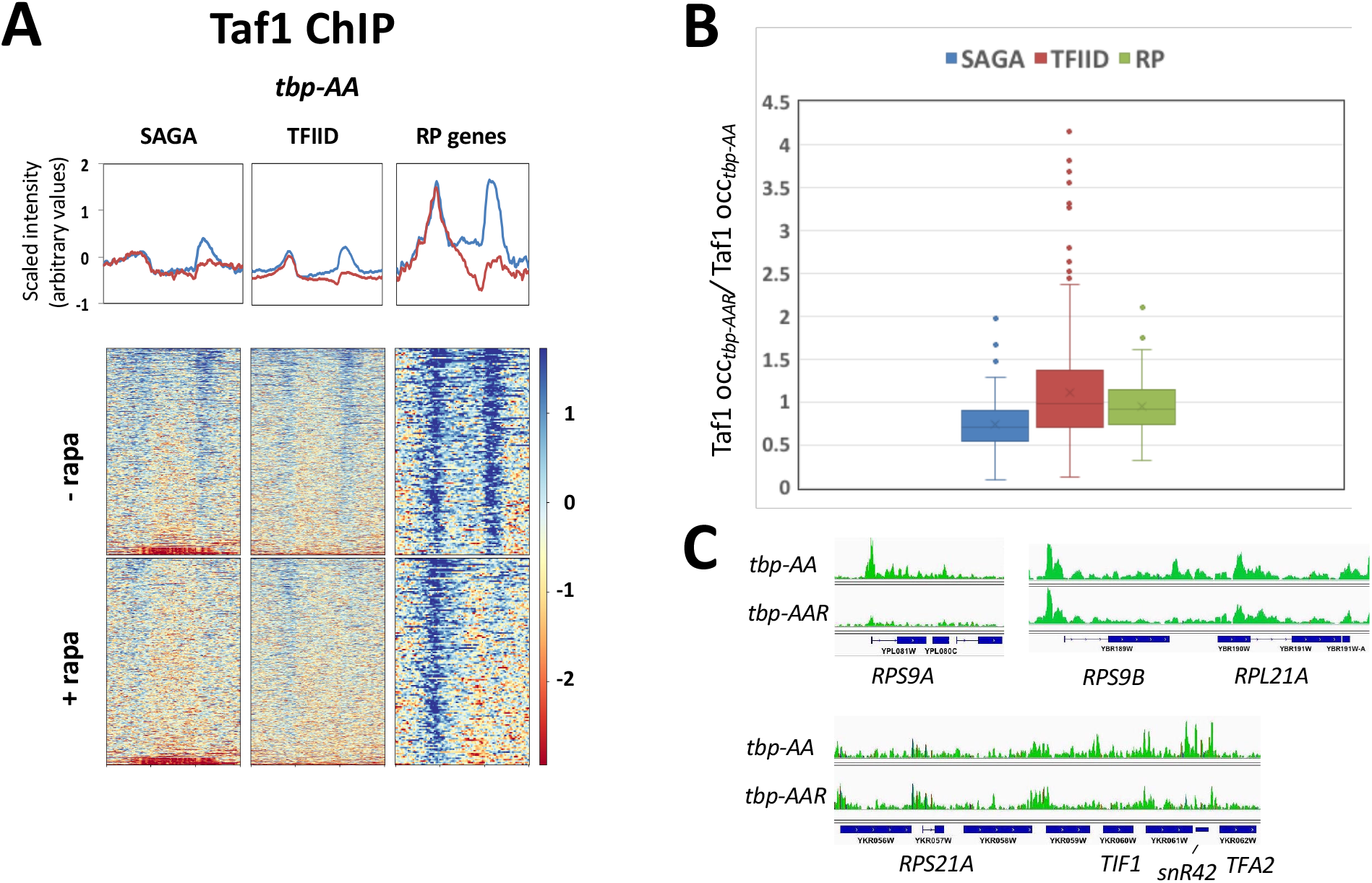
Depletion of TBP has little effect on Taf1 occupancy. (A) Normalized Taf1 occupancy in *tbp-AA* yeast, without and with 1 hr rapamycin treatment, mapped to all SAGA-dominated, non-RP TFIID-dominated, and RP genes. Genes were normalized for length, aligned by coding sequence (CDS) start and stop, and sorted according to average signal intensity. The 22 SAGA-dominated genes at the bottom of the heat maps were removed before calculating averages used in the line graphs, as these were almost Ty1 elements that had higher intensity in the input control than in the Taf1 ChIP sample, and therefore yielded negative values in the heat map. (B) Ratios of Taf1 occupancy in *tbp-AA* yeast in the presence and absence of rapamycin are shown in box and whisker plots, as in Figure 2, for the ~1000 genes having highest occupancy by Pol II (see Methods), sorted into SAGA-dominated, non-RP TFIID-dominated, and RP genes. (C) Browser scans showing normalized Taf1 occupancy in *tbp-AA* yeast with and without rapamycin treatment.

### Variation in TBP/Taf1 ratio at gene promoters

Although essentially all genes transcribed by Pol II in yeast depend on both TFIID and the SAGA complex for normal levels of transcription, differential effects of conditional mutations in SAGA and TFIID components led to categorization of about 10% of yeast genes as SAGA-dominated and about 90% as being TFIID-dominated [4, 8]. This categorization was supported by subsequent genome-wide ChIP results showing higher levels of TFIID components at TFIID-dominated than at SAGA-dominated genes [29, 30]. However, a more recent study using ChEC-seq found comparable levels of Taf1 association at SAGA-dominated and TFIID-dominated promoters [13], while two other reports showed that depletion of TFIID and SAGA components resulted in decreased Pol II association equally at SAGA- and TFIID-dominated genes; effects at genes having consensus TATA elements and those lacking consensus TATA elements were also indistinguishable when examined across all genes [7, 9].

To gain further insight into whether these gene categories reflect mechanistic differences, we compared the ratio of TBP and Taf1 occupancy for the 1000 genes having highest Pol II occupancy (normalized for gene length) as determined in previous ChIP-seq experiments [16]. After removing genes showing anomalous ChIP peaks or being proximate to tRNA genes (see Methods), the remaining cohort included 154 SAGA-dominated genes, 534 TFIID-dominated genes, and 136 RP genes that were considered separately from SAGA- or TFIID-dominated genes [4].

Plotting TBP occupancy against Taf1 occupancy for SAGA-dominated, TFIID-dominated, and RP genes revealed strong correlations for all three groups and slopes that differed considerably (Fig. 6A). (Note that the slopes do not reflect molar ratio, as it is not possible to determine this directly from ChIP experiments using distinct antibodies for Taf1 and TBP.) RP genes clearly formed a distinct cohort and exhibited the lowest TBP/Taf1 ratios, consistent with previous studies indicating high TFIID occupancy and dependence of these genes (Fig. 6A-B) [5, 6, 31]. SAGA-dominated genes showed the lowest levels of Taf1 occupancy relative to TBP; TFIID-dominated genes were intermediate in their TBP/Taf1 ratios between SAGA-dominated and RP genes and significantly different as a class from both (respective medians were 3.2, 1.9, and 1.4; p-values < 1 × 10^−5^ for all pairwise comparisons using Mann-Whitney U-test) (Fig. 6B).

**Figure 6.**
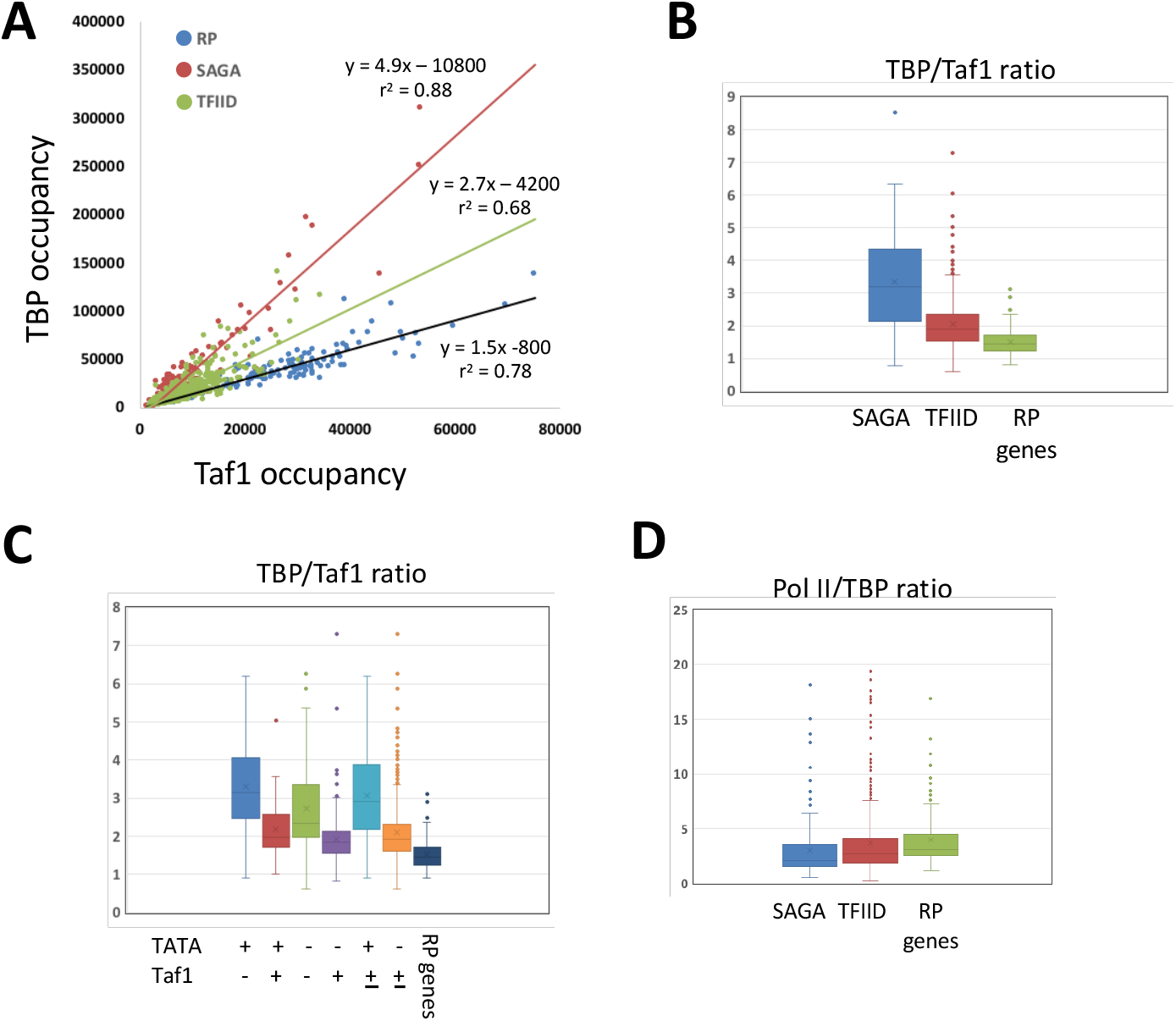
Differential TBP/Taf1 ratios at SAGA- and TFIID-dominated genes, and at gene promoters having and lacking consensus TATA elements. (A) Normalized occupancy of TBP plotted against Taf1 occupancy for the ~1000 genes (see Methods) most highly occupied by Pol II, separated into SAGA-dominated, non-RP TFIID-dominated, and RP genes. (B) Ratios of TBP to Taf1 occupancy in *kin28-AA* yeast in the absence of rapamycin are shown in box and whisker plots, as in Figure 2, for the ~1000 genes having highest occupancy by Pol II (see Methods), sorted into SAGA-dominated, non-RP TFIID-dominated, and RP genes. (C) As in (B), with genes sorted by the presence or absence of a consensus TATA element and enrichment or depletion of Taf1 [29]. RP genes were removed from these categories and are shown separately. (D) Ratios of Pol II, normalized for gene length, to TBP occupancy in *kin28-AA* yeast in the absence of rapamycin are shown in box and whisker plots, as in Figure 2, for the ~1000 genes having highest occupancy by Pol II (see Methods), sorted into SAGA-dominated, non-RP TFIID-dominated, and RP genes.

RP genes are categorized by function; the high Taf1 levels and low TBP/Taf1 ratios observed for this cohort therefore reflect properties independent of their categorization that likely reflect mechanistically distinct behavior. In contrast, the disparate TBP/Taf1 ratios observed for SAGA-dominated and TFIID-dominated genes corroborate their categorization but do not provide evidence for distinct mechanisms of transcriptional activation. Rhee and Pugh measured occupancy of PIC components including Taf1 and TBP using ChIP-exo, and re-categorized yeast genes as enriched or depleted for Taf1, and containing or lacking a consensus TATA element [29]. We plotted TBP/Taf1 ratios for these groups and noted that TATA-containing genes behaved distinctly (again with overlap) from genes lacking a consensus TATA element (Fig. 6C). Since the property of having or lacking a TATA element is independent of measurements of Taf1 and TBP occupancy, this provides additional evidence for distinct mechanisms of PIC assembly at these two gene categories. Finally, we note that the ratio of Pol II (normalized for gene length) to TBP occupancy displays a trend that is the opposite of that seen for TBP/Taf1 ratios (respective medians for SAGA-dominated, TFIID-dominated, and RP genes of 2.1, 2.7, and 3.1), suggesting that TFIID may be more effective than SAGA at facilitating Pol II recruitment.

## Discussion

A principal conclusion of this work is that depletion or inactivation of Kin28 results in increased occupancy, as measured by ChIP, of Taf1 and Taf4 at promoters genome-wide. This conclusion presumes that ChIP signal accurately reflects occupancy. An alternative possibiliity is that conformational changes affect the efficiency of immunoprecipitation, and that inhibition of Pol II promoter escape by depletion or inactivation of Kin28 locks TFIID in a configuration in which ChIP of Taf subunits is more efficient than in the presence of Kin28. Consistent with this notion, human TFIID undergoes major conformational changes upon binding of TFIID (together with TBP) to promoter DNA and TFIIA [11, 32]. However, Taf1 and Taf4 are situated in distinct regions of TFIID: TFIID comprises three lobes, with Taf4 situated at the apical ends of the two outer lobes in the promoter-unbound state while Taf1 is on the opposite side of the central lobe. It thus seems unlikely that conformational changes would have the same effect on accessibility of these two Tafs, while also having negligible effect on TBP accessibility. We therefore interpret our results as indicating that occupancy of Taf1 and Taf4, and likely TFIID as a unit, is stabilized when Pol II escape is inhibited by depletion or inactivation of Kin28.

Mediator occupancy at gene promoters is stabilized by depletion or inactivation of Kin28, and also in the absence of the Pol II CTD, which is targeted for phosphorylation by Kin28 [15, 18]. This stabilization is suppressed by depletion of Pol II or TBP, which results in a severe reduction in Mediator occupancy at proximal promoter regions and, in the case of TBP depletion, an upstream shift of Mediator ChIP signal to the UAS regions that are sites of initial Mediator recruitment by gene-specific activators [21]. Here, we show that increased Taf1 occupancy caused by Kin28 depletion is suppressed by simultaneous depletion of TBP. In the case of Mediator, decreased occupancy caused by depletion of Pol II or TBP likely reflects loss of direct interactions between Mediator and PIC components, including Pol II [21, 33]. TFIID occupancy may similarly depend in part on interactions with other PIC components, and the decreased occupancy seen upon simultaneous depletion of TBP and Kin28 compared to Kin28 alone could reflect loss of interactions with TBP or with PIC components or other factors whose recruitment depends on TBP. Alternatively, TFIID binding to promoter regions may be destabilized by clashes with GTFs that bind subsequent to TFIID [11]; inhibition of Pol II promoter escape could interfere with later steps in transcription initiation, thereby mitigating potential clashes with TFIID.

Interestingly, when Kin28 is present, depletion of TBP had little effect on Taf1 occupancy. The lack of effect on Taf1 occupancy of depletion of TBP indicates that recruitment of Tafs in TFIID does not depend much on TBP, consistent with previous reports [3, 6]. Given that TBP depletion interrupts the normal transcription cycle, one might have predicted increased Taf occupancy to accompany TBP depletion, analogous to interruption of the transcription cycle by depletion of Kin28. The fact that no such increase in Taf1 occupancy was observed indicates that TBP, or PIC components or other factors whose recruitment depends on TBP, are required for such elevated Taf occupancy, consistent with TBP being required for elevated Taf1 occupancy caused by Kin28 depletion.

Unlike Mediator and TFIID, TBP occupancy measured by ChIP does not increase upon depletion of Kin28 (Fig. 3). Thus, TBP occupancy does not strictly correlate with binding of Tafs through the transcription cycle. This does not mean that Tafs are not required for recruitment of TBP to promoters, as it may be that once TBP has been recruited, interactions with promoter DNA and other PIC components are sufficient to allow continued TBP occupancy. However, our findings do suggest that promoter occupancy by TBP and Tafs are uncoupled during the normal transcription cycle, with Taf occupancy being transient and TBP occupancy being stable. Such uncoupling has been suggested during transcription of metazoan genes based on structural and biochemical studies of TFIID [11].

We also took advantage of our ChIP-seq data to examine TBP/Taf ratios at promoters across the yeast genome. Our data corroborate previous work showing increased Taf occupancy, relative to TBP, at TFIID-dominated promoters relative to SAGA-dominated promoters (Fig. 6) [3–6, 8, 30]. We also found a marked difference in TBP/Taf1 occupancy ratios at promoters having and lacking consensus TATA elements (Mann Whitney U Test z-score -12.5 for TATA+ vs TATA-promoters; p < 10^−5^; Fig. 6), providing quantitative support for previous work [29]. Because the presence or absence of a consensus TATA element is a property completely independent of occupancy by Tafs or TBP, this difference strongly suggests mechanistic differences between these two categories of promoters. At the same time, there is considerable overlap between the TBP/Taf ratios (and of other properties) between TATA+ and TATA-promoters; the most parsimonious explanation is that either of at least two distinct mechanisms or pathways can operate at both types of promoters, but their relative efficiency differs depending on the presence of a consensus TATA element. This notion is consistent with the idea that the categorization of genes as SAGA- and TFIID-dominated reflects a continuum in terms of regulation, not a rigid dichotomy [4, 5, 34]. We also observed a greater increase in occupancy by Taf1 and Taf4 upon Kin28 depletion at TFIID-dominated than at SAGA-dominated promoters (Fig. 2), and at TATA-than at TATA+ promoters (not shown). This may simply reflect the higher occupancy at TFIID-dominated/TATA-promoters seen in the presence of Kin28, or it may be that Tafs are present in more than one configuration which differ in their response to Kin28 depletion and which vary in proportion at the two categories of promoters.

## Materials and Methods

### Yeast strains and growth

Yeast strains used in this study are listed in Table S1. Cultures were grown in CSM-ura (0.67% yeast nitrogen base without amino acids and 2% glucose supplemented with CSM-ura dropout mix (Bio101)) (Figure 1) or yeast peptone dextrose (YPD) media (1% bacto-yeast extract, 2% bacto-peptone extract, 2% glucose) at 30°C with shaking at 100 rpm. For experiments using the *kin28-as* analog sensitive mutant, 1-Naphythyl-PP1 (NaPP1) was added to 1 μg/ml and incubation continued for 30 min before cross-linking. For experiments using anchor away strains, rapamycin (LC Laboratories, Woburn, MA) was added one hour prior to crosslinking to a final concentration of 1 μg/mL from a 1 mg/mL stock, stored in ethanol at - 20°C for not more than one month. (Concentration of rapamycin stock solutions was determined using A_267_ = 42 and A_277_ = 54 for a 1 mg/ml solution.)

### ChIP and ChIP-seq

Whole cell extracts (WCE) were prepared from 50 mL cultures as described previously, yielding 600-800 μl of WCE [16]. Immunoprecipitations were performed using 180 μl of WCE for analysis by qPCR or the entire WCE less 36 μl saved as “input” for ChIP-seq. Samples were incubated overnight at 4°C with 2.5-5 μg anti-TBP (58C9, Abcam, or 5 μL serum, generous gift from A. Weil, Vanderbilt University) or 2.0 μL anti-Taf1 or anti-Taf4 (serum, generous gift from J. Reese and Song Tan, Penn State University). Immunoprecipitated DNA was purified using 30 μl of protein A beads (Sigma), which were washed prior to DNA elution and cross-link reversal as previously described [16, 35].

Analysis of ChIP samples by qPCR was performed on an Applied Biosystems StepOnePlus instrument, using SYBR Green master mix and ROX passive dye (ThermoFisher/USB/Affymetrix). Each reaction contained 0.5 μl of IP DNA or of a 1:100 dilution of input DNA in a 12.5 μl volume, and was performed in duplicate. Error bars in Figure 1 represent standard deviation based on biological replicates. IP samples were normalized against input, and then against IP/input values for *SNR6* (a Pol III transcribed gene) or a nontranscribed region of ChrV [36]. Oligonucleotides used for qPCR are shown in Table S2.

Library preparation for Illumina paired-end sequencing was performed with the NEBNext Ultra II library preparation kit (New England Biolabs) according to manufacturer’s protocol and barcoded using NEXTflex barcodes (BIOO Scientific, Austin, TX) or NEBNext Multiplex Oligos for Illumina. In some experiments, a size selection step was performed on barcoded libraries by isolating fragment sizes between 200-500 bp on a 2% E-Gel EX agarose gel apparatus (ThermoFisher Scientific). Sequencing was performed on the Illumina NextSeq 500 platform at the University of Buffalo next-generation sequencing and expression analysis core (University of Buffalo, State University of New York, Buffalo, NY) or at the Illumina NextSeq platform at the Wadsworth Center, New York State Department of Health (Albany, NY).

### ChIP-Seq analysis

Unfiltered sequencing reads were aligned to the *S. cerevisiae* reference genome (Saccer3) using bwa [37]. Up to 1 mismatch was allowed for each aligned read. Reads mapping to multiple sites were retained to allow evaluation of associations with non-unique sequences [37] and duplicate reads were retained. Calculation of coverage, comparisons between different data sets, and identification of overlapping binding regions were preceded by library size normalization, and were performed with the “chipseq” and “GenomicRanges” packages in BioConductor [38]. Alternatively, reads were aligned and analysis conducted using the Galaxy platform [39] and Excel. For metagene analysis, including heat maps, reads were normalized against input from *kin28*-AA yeast grown in the absence of rapamycin (KHW127). Taf1 and TBP occupancy were determined as read depth over the 300 bp upstream of coding sequence using BedCov in SamTools [40]. Ten genes exhibiting anomalous Taf1/Taf4 signal were removed prior to further analysis; the comprised 5 genes present in the rDNA locus and five additional genes displaying anomalous peaks. Genes proximate to tRNA genes (within 500 bp upstream of the 5’ end of the ORF) were removed in analyses of TBP occupancy. For analysis of Pol II/TBP ratios, 14 genes having ratios > 20 were removed from consideration; these comprised nine dubious ORFs, a Ty element, *STE2* (which is expressed in the *Mat a* strain used for Pol II ChIP-seq but not in the *Mat α kin28-AA* strain), *CSS1, RPS29A* and *RPS29B*. The 1000 genes having highest Pol II occupancy were obtained using BedCov to obtain read depth over coding sequences using Pol II ChIP-seq data from [16] (strain BY4741 grown at 30°C in YPD medium). Genes designated as SAGA-dominated and TFIID-dominated were obtained from [4], and genes designated as containing or not containing a consensus TATA element, and being Taf1-enriched or Taf1-depleted, were obtained from [29]. Occupancy profiles were normalized for read depth and generated using the Integrative Genomics Viewer [41]. Gene ontology analysis was performed using the Generic Gene Ontology Term Finder (https://go.princeton.edu/cgi-bin/GOTermFinder/GOTermFinder) [27]. Hypergeometric test p-values were calculated using the online calculator at http://www.alewand.de/stattab/tabdiske.htm, and the Mann Whitney U Test was performed using the online calculators at http://astatsa.com/WilcoxonTest/ and https://www.socscistatistics.com/tests/mannwhitney/default2.aspx.

### Data deposition

ChIP-seq reads have been deposited in the NCBI Short Read Archive under project number PRJNA541522. We also used previously published ChIP-seq data deposited at the NCBI Short Read Archive under accession numbers SRP047524 [16] and PRJNA413080 [21].

## Acknowledgements

This work was supported by grant MCB1516839 to RHM, and was also funded in part by the NIH Intramural Research Program at the National Library of Medicine (ZIZ and DL). We thank Tony Weil (Vanderbilt University) and Joe Reese and Song Tan (Pennsylvania State University) for generous gifts of antibody sera, and Chhabi Govind and Kevin Struhl for yeast strains and reagents. We also acknowledge Joe Wade for helpful discussions and Mike Palumbo for advice on using SamTools. We gratefully acknowledge help from the Wadsworth Center Applied Genomics Technology and Tissue Culture and Media Cores.

---

**Supplementary Table 1.**
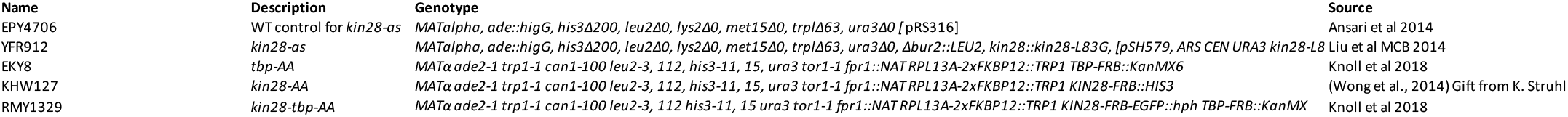
Yeast strains used in this study.

**Supplementary Table 2.**
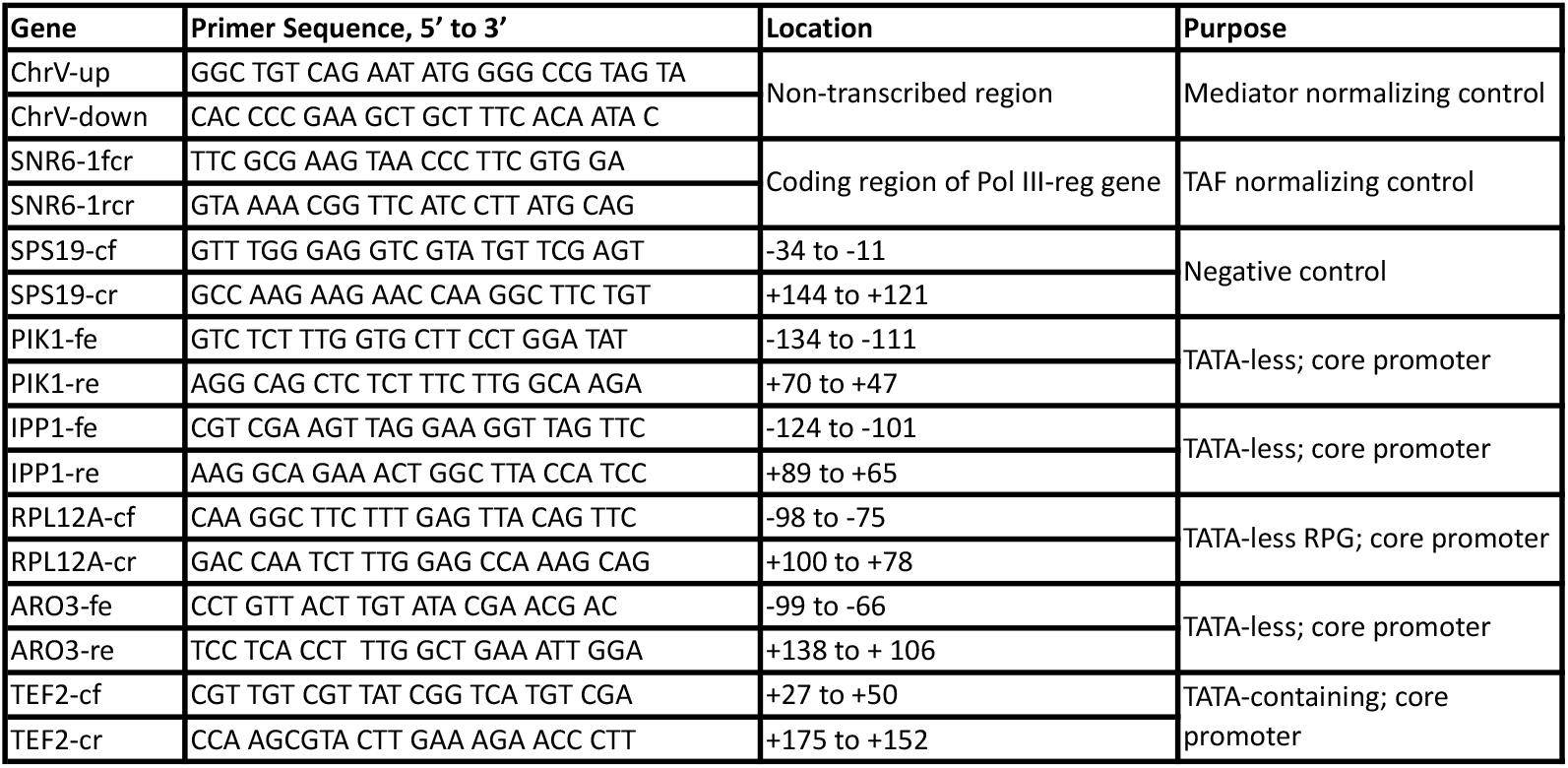
Oligonucleotides used in this study.

## References

1. Sainsbury S, Bernecky C, Cramer P. Structural basis of transcription initiation by RNA polymerase II. Nat Rev Mol Cell Biol. 2015;16(3):129–43.

2. Eisenmann DM, Arndt KM, Ricupero SL, Rooney JW, Winston F. SPT3 interacts with TFIID to allow normal transcription in Saccharomyces cerevisiae. Genes Dev. 1992;6(7):1319–31.

3. Shen WC, Bhaumik SR, Causton HC, Simon I, Zhu X, Jennings EG, et al. Systematic analysis of essential yeast TAFs in genome-wide transcription and preinitiation complex assembly. EMBO J. 2003;22(13):3395–402.

4. Huisinga KL, Pugh BF. A genome-wide housekeeping role for TFIID and a highly regulated stress-related role for SAGA in Saccharomyces cerevisiae. Mol Cell. 2004;13(4):573–85.

5. Kuras L, Kosa P, Mencia M, Struhl K. TAF-Containing and TAF-independent forms of transcriptionally active TBP in vivo. Science. 2000;288(5469):1244–8.

6. Li XY, Bhaumik SR, Green MR. Distinct classes of yeast promoters revealed by differential TAF recruitment. Science. 2000;288(5469):1242–4.

7. Baptista T, Grunberg S, Minoungou N, Koster MJE, Timmers HTM, Hahn S, et al. SAGA Is a General Cofactor for RNA Polymerase II Transcription. Mol Cell. 2017;68(1):130–43 e5.

8. Lee TI, Causton HC, Holstege FC, Shen WC, Hannett N, Jennings EG, et al. Redundant roles for the TFIID and SAGA complexes in global transcription. Nature. 2000;405(6787):701–4.

9. Warfield L, Ramachandran S, Baptista T, Devys D, Tora L, Hahn S. Transcription of Nearly All Yeast RNA Polymerase II-Transcribed Genes Is Dependent on Transcription Factor TFIID. Mol Cell. 2017;68(1):118–29 e5.

10. Joo YJ, Ficarro SB, Soares LM, Chun Y, Marto JA, Buratowski S. Downstream promoter interactions of TFIID TAFs facilitate transcription reinitiation. Genes Dev. 2017;31(21):2162–74.

11. Patel AB, Louder RK, Greber BJ, Grunberg S, Luo J, Fang J, et al. Structure of human TFIID and mechanism of TBP loading onto promoter DNA. Science. 2018;362(6421).

12. Jeronimo C, Robert F. The Mediator Complex: At the Nexus of RNA Polymerase II Transcription. Trends Cell Biol. 2017;27(10):765–83.

13. Grunberg S, Henikoff S, Hahn S, Zentner GE. Mediator binding to UASs is broadly uncoupled from transcription and cooperative with TFIID recruitment to promoters. EMBO J. 2016;35(22):2435–46.

14. Jeronimo C, Langelier MF, Bataille AR, Pascal JM, Pugh BF, Robert F. Tail and Kinase Modules Differently Regulate Core Mediator Recruitment and Function In Vivo. Mol Cell. 2016;64(3):455–66.

15. Jeronimo C, Robert F. Kin28 regulates the transient association of Mediator with core promoters. Nat Struct Mol Biol. 2014;21(5):449–55.

16. Paul E, Zhu ZI, Landsman D, Morse RH. Genome-wide association of mediator and RNA polymerase II in wild-type and mediator mutant yeast. Mol Cell Biol. 2015;35(1):331–42.

17. Petrenko N, Jin Y, Wong KH, Struhl K. Mediator Undergoes a Compositional Change during Transcriptional Activation. Mol Cell. 2016;64(3):443–54.

18. Wong KH, Jin Y, Struhl K. TFIIH phosphorylation of the Pol II CTD stimulates mediator dissociation from the preinitiation complex and promoter escape. Mol Cell. 2014;54(4):601–12.

19. Fan X, Chou DM, Struhl K. Activator-specific recruitment of Mediator in vivo. Nat Struct Mol Biol. 2006;13(2):117–20.

20. Fan X, Struhl K. Where does mediator bind in vivo? PLoS One. 2009;4(4):e5029.

21. Knoll ER, Zhu ZI, Sarkar D, Landsman D, Morse RH. Role of the pre-initiation complex in Mediator recruitment and dynamics. Elife. 2018;7.

22. Buratowski S. Progression through the RNA polymerase II CTD cycle. Mol Cell. 2009;36(4):541–6.

23. Liu Y, Kung C, Fishburn J, Ansari AZ, Shokat KM, Hahn S. Two cyclin-dependent kinases promote RNA polymerase II transcription and formation of the scaffold complex. Mol Cell Biol. 2004;24(4):1721–35.

24. Bataille AR, Jeronimo C, Jacques PE, Laramee L, Fortin ME, Forest A, et al. A universal RNA polymerase II CTD cycle is orchestrated by complex interplays between kinase, phosphatase, and isomerase enzymes along genes. Mol Cell. 2012;45(2): 158–70.

25. Haruki H, Nishikawa J, Laemmli UK. The anchor-away technique: rapid, conditional establishment of yeast mutant phenotypes. Mol Cell. 2008;31(6):925–32.

26. Teytelman L, Thurtle DM, Rine J, van Oudenaarden A. Highly expressed loci are vulnerable to misleading ChIP localization of multiple unrelated proteins. Proc Natl Acad Sci U S A. 2013;110(46):18602–7.

27. Boyle EI, Weng S, Gollub J, Jin H, Botstein D, Cherry JM, et al. GO::TermFinder--open source software for accessing Gene Ontology information and finding significantly enriched Gene Ontology terms associated with a list of genes. Bioinformatics. 2004;20(18):3710–5.

28. Harbison CT, Gordon DB, Lee TI, Rinaldi NJ, Macisaac KD, Danford TW, et al. Transcriptional regulatory code of a eukaryotic genome. Nature. 2004;431(7004):99–104.

29. Rhee HS, Pugh BF. Genome-wide structure and organization of eukaryotic pre-initiation complexes. Nature. 2012;483(7389):295–301.

30. Venters BJ, Wachi S, Mavrich TN, Andersen BE, Jena P, Sinnamon AJ, et al. A comprehensive genomic binding map of gene and chromatin regulatory proteins in Saccharomyces. Mol Cell. 2011;41(4):480–92.

31. Mencia M, Moqtaderi Z, Geisberg JV, Kuras L, Struhl K. Activator-specific recruitment of TFIID and regulation of ribosomal protein genes in yeast. Mol Cell. 2002;9(4):823–33.

32. Cianfrocco MA, Kassavetis GA, Grob P, Fang J, Juven-Gershon T, Kadonaga JT, et al. Human TFIID binds to core promoter DNA in a reorganized structural state. Cell. 2013; 152(1-2):120–31.

33. Soutourina J, Wydau S, Ambroise Y, Boschiero C, Werner M. Direct interaction of RNA polymerase II and mediator required for transcription in vivo. Science. 2011;331(6023):1451–4.

34. Petrenko N, Jin Y, Dong L, Wong KH, Struhl K. Requirements for RNA polymerase II preinitiation complex formation in vivo. Elife. 2019;8.

35. Kuo MH, Allis CD. In Vivo Cross-Linking and Immunoprecipitation for Studying Dynamic Protein:DNA Associations in a Chromatin Environment. Methods. 1999;19(3):425–33.

36. Keogh MC, Buratowski S. Using chromatin immunoprecipitation to map cotranscriptional mRNA processing in Saccharomyces cerevisiae. Methods Mol Biol. 2004;257:1–16.

37. Seoighe C, Wolfe KH. Updated map of duplicated regions in the yeast genome. Gene. 1999;238(1):253–61.

38. Gentleman RC, Carey VJ, Bates DM, Bolstad B, Dettling M, Dudoit S, et al. Bioconductor: open software development for computational biology and bioinformatics. Genome Biol. 2004;5(10):R80.

39. Goecks J, Nekrutenko A, Taylor J. Galaxy: a comprehensive approach for supporting accessible, reproducible, and transparent computational research in the life sciences. Genome Biol. 2010;11(8):R86.

40. Li H, Handsaker B, Wysoker A, Fennell T, Ruan J, Homer N, et al. The Sequence Alignment/Map format and SAMtools. Bioinformatics. 2009;25(16):2078–9.

41. Robinson JT, Thorvaldsdottir H, Winckler W, Guttman M, Lander ES, Getz G, et al. Integrative genomics viewer. Nat Biotechnol. 2011;29(1):24–6.

